# Multi-omics Analysis of Aging Liver Reveals Changes in Endoplasmic Stress and Degradation Pathways in Female Nonhuman Primates

**DOI:** 10.1101/2023.08.21.554149

**Authors:** Sobha Puppala, Jeannie Chan, Kip D. Zimmerman, Zeeshan Hamid, Isaac Ampong, Hillary F. Huber, Ge Li, Avinash Y. L. Jadhav, Cun Li, Peter W. Nathanielsz, Michael Olivier, Laura A. Cox

## Abstract

The liver is critical for functions that support metabolism, immunity, digestion, detoxification, and vitamin storage. Aging is associated with severity and poor prognosis of various liver diseases such as nonalcoholic fatty liver disease (NAFLD). Previous studies have used multi-omic approaches to study liver diseases or to examine the effects of aging on the liver. However, to date, no studies have used an integrated omics approach to investigate aging-associated molecular changes in the livers of healthy female nonhuman primates. The goal of this study was to identify molecular changes associated with healthy aging in the livers of female baboons (*Papio* sp., n=35) by integrating multiple omics data types (transcriptomics, proteomics, metabolomics) from samples across the adult age span. To integrate omics data, we performed unbiased weighted gene co-expression network analysis (WGCNA), and the results revealed 3 modules containing 3,149 genes and 33 proteins were positively correlated with age, and 2 modules containing 37 genes and 216 proteins were negatively correlated with age. Pathway enrichment analysis showed that unfolded protein response (UPR) and endoplasmic reticulum (ER) stress were positively associated with age, whereas xenobiotic metabolism and melatonin and serotonin degradation pathways were negatively associated with age. The findings of our study suggest that UPR and a reduction in reactive oxygen species generated from serotonin degradation could protect the liver from oxidative stress during the aging process in healthy female baboons.

## Introduction

Aging is a key risk factor influencing many diseases, including liver diseases (Radonjic et al 2022, Jin et al 2020). The liver is an important metabolic organ that is essential for maintaining whole body homeostasis via regulation of energy metabolism, xenobiotic and endobiotic clearance, and biosynthesis of lipids and proteins (Hunt et al 2019). Recent studies show that genetic and epigenetic modifications from aging leads to a liver with cellular senescence and low-grade inflammation, in which mitochondrial function and nutrient sensing pathways are the underlying mechanisms (Yao et al 2023, Hunt et al 2019). However, the molecular changes that accompany aging in healthy liver have not been thoroughly investigated.

Previous studies have primarily used rodents to understand the aging-related alterations in liver (Zhao et al 2022). For example, Ori et al (2015) and Smiljanic et al (2013) identified decline in gene expression, translation, protein abundance, and phosphorylation in livers of old (24-month-old) rats. Capel et al found aging-related genes related to xenobiotic and fatty acid metabolism, retinoid X receptor function, and oxidative stress in the aged mouse liver. Chishti et al (2013) found down-regulation of many genes involved in lipid metabolism and cell growth in livers of aged rats. Musicco et al (2011) identified differentially expressed hepatic mitochondrial proteins between old (28-month-old) and young adult (12-month-old) rats. Honma et al (2011 and 2012) found age-dependent changes in lipid and carbohydrate metabolism in a senescence-accelerated mouse model. Pagliassotti et al (2017) identified reduction of several metabolites in livers of aged mice. The above studies on aging rodent models showed changes in liver, but fundamental differences in the aging process between rodents and humans have hindered direct translation of findings in rodents to humans (Zhao et al 2022, Colman 2018, Messaudi and Ingram 2012). Previous studies in nonhuman primates (NHP) and humans have focused on dysregulated processes in the aging liver related to nonalcoholic fatty liver disease (NAFLD), nonalcoholic steatohepatitis (NASH), and fibrosis (Li et al 2022, Xu et al 2022). NHPs are ideal translational models because they share strikingly similar genetic, physiological, and behavior traits with humans (Tarantal et al 2022). Our recent NHP studies aim to better understand human disease processes have focused on genetics, epigenetics, caloric restriction, maternal caloric restriction and offspring health, maternal obesity and offspring health, NASH, and steatosis (Zimmerman et al 2023, Puppala et al 2022 and 2018, Cox et al 2017). In humans, studies comparing diseased vs. healthy liver have shown decline in cellular and biological functions (Wang et al 2023, Campagnoli et al 2022). However, little is known about age-related molecular changes in healthy liver.

The goal of this study was to use an integrated omics method to analyze transcriptomic, proteomic, and metabolomic data together to determine molecular changes associated with healthy aging in liver. The present study included 35 female baboons (*Papio* sp.) across the adult age span. In this model of healthy primate aging, we found unfolded protein response (UPR) and endoplasmic reticulum (ER) stress were associated with age. We also found age-associated changes in xenobiotic metabolism and melatonin and serotonin degradation. Our results provide insights into the molecular changes that accompany the aging process to maintain the liver in a healthy condition.

## Materials and Methods

### Animals and liver necropsy collection

The study included 35 females ranging in age from 6.0 to 22.1 years (y) (human equivalent ∼24 to 88 y), median age 13.2 y (human equivalent ∼50 y, Bronikowski et al 2002). Sample collection procedures from baboons were described in a recent paper (Cox et al 2022). Following cardiac asystole, liver tissue samples were collected within an hour from time of death between 8:00-10:00 AM to minimize potential variation from circadian rhythms, snap-frozen in liquid nitrogen, and stored at -80°C until use (Yang et al 2017).

### Morphometric and clinical measures

Morphometrics were measured prior to necropsy using standard anatomical landmarks as described previously (Li et al 2017, Chavez et al 2009). Details of blood sample collection, timings of collection, and details of assay precision for all measures were published (Li et al 2017). The morphometric and clinical measures used for our study are in Supplemental Tables ST1. A Pearson correlation was performed to determine if there was any significant association between age or age-squared with morphometric and clinical traits.

### Ethics statement, study design, and data collection

All animal procedures took place at the Southwest National Primate Research Center (SNPRC) at Texas Biomedical Research Institute (Texas Biomed), were approved by the Texas Biomed Animal Care and Use Committee (IACUC), and were conducted in facilities approved by the Association for Assessment and Accreditation of Laboratory Animal Care (AAALC). The Texas Biomed animal use programs operate according to all National Institutes of Health (NIH) and U.S. Department of Agriculture guidelines and are directed by board certified veterinarians. Details of animal care and housing were published (see Cox et al 2022). Animals were maintained on *ad libitum* normal monkey chow diet (Monkey Diet 5LEO, Purina, St Louis, MO, USA) throughout their lifespan and water was continuously available along with complete veterinary care by SNPRC veterinary staff throughout their lives (Cox et al 2022). Fresh serum samples were collected under sedation (ketamine 10 mg/kg IM) from overnight fasted animals and stored in aliquots at -80°C until analysis. After study completion, baboons were euthanized by exsanguination under general anesthesia (initiated with ketamine 10 mg/kg and maintained with isoflurane 1-2% INH) by a SNPRC veterinarian in accordance with American Veterinary Medical Association (AVMA) guidelines (Leary et al 2013). Humane endpoint was confirmed by the Texas Biomed IACUC protocol as lack of pulse, breathing, corneal reflex, and response to firm toe pinch, and inability of the veterinarian to hear respiratory sounds and heartbeat by stethoscope (Mahaney et al 2018). Following cardiac asystole, all tissues were collected according to the same protocol.

### Transcriptomics

#### RNA Isolation

Approximately 5 mg of frozen liver was homogenized in 1 ml RLT buffer (Qiagen) using a BeadBeater (BioSpec) with zirconia/silica beads, and RNA was extracted using the Zymo Direct-zol RNA Miniprep Plus according to manufacturer’s instructions. RNA quality was assessed by TapeStation high sensitivity RNA ScreenTape (Agilent). RNA was stored at −80°C.

#### Sequence Data Generation

cDNA libraries were generated from RNA samples by using Kapa RNA HyperPrep Kit with RiboErase kit with quality assessed by Agilent High Sensitivity D1000 ScreenTape according to the manufacturers’ protocols. cDNA libraries were pooled and sequenced using the Illumina Flow cell v1.0 reagent kit for 2x150 paired-end reads on an NovaSeq 6000 Sequencer (Cox et 2022).

#### Sequence Data Analysis

Low quality bases with Phred scores below 30 were removed prior to alignment. Trimmed reads were aligned against the olive baboon reference (Panu_3.0, GCF_000264685.3) using HISAT2 (Kim, Langmead, and Salzberg 2015). Aligned reads were quantified using an expectation-maximization algorithm (Xing et al 2006) with Panu_3.0 annotation (NCBI release 103). Transcripts (without read counts) across all samples were filtered out and then normalized by the trimmed mean of M values method (Robinson and Oshlack 2010). Raw read counts were filtered to remove those with less than or equal to 30 counts across all samples resulting in 25,562 transcripts that passed quality filters.

### Proteomics

#### Sample Processing

Details of preparation of liver proteomic samples were as described (Hamid et al 2022, Cox et al 2022). Approximately 5 mg of each liver tissue sample was homogenized in Tris buffer, precipitated overnight in acetone at −20°C, and centrifuged at 12,000 g for 10 min. The protein pellet was dried, reconstituted in 100 mM of ammonium bicarbonate and quantified. One hundred mg of protein was reduced for 30 min in dark, and digested overnight with trypsin. Samples were cleaned and desalted using Thermo Scientific Pierce C18 Tips, dried, and reconstituted in 0.1% formic acid.

#### LC/MS Data Acquisition and Analysis

LC/MS data were acquired as described (Cox et al 2022, Hamid et al 2022). One mg of each sample was loaded on a PepMap RSLC C18 easy-spray column using Easy-nLC 1200 coupled to an Orbitrap Lumos Tribrid Mass Spectrometer (Thermo Scientific). Peptides were separated using a 3 hr gradient of Mobile phase A and Mobile Phase B. Peptides were eluted according to the gradient program. Mass spectrometer data were acquired in MS1 scan mode. All data acquisition was done using Thermo Scientific Xcalibur software. MS raw data were analyzed using MetaMorpheus (Hamid et al 2022, Miller et al 2019) and the *P. anubis* reference proteome database from Uniprot with 44,721 entries (UP000028761). Peptide and protein quantification were done using the FlashLFQ approach. Protein intensities were normalized using global intensity normalization. In the final normalized data missing values were imputed using random forest imputation workflow (Hamid et al 2022, Stekhoven and Buhlmann 2012).

### Metabolomics

#### Sample Processing

Extraction of metabolites from liver samples was performed following a protocol adopted from a previously described study (Misra et al 2019). Briefly, aliquots (15 μL) of liver homogenates were subjected to sequential solvent extraction, The extracts were then dried under vacuum at 4°C prior to chemical derivatization (silylation reactions). Tubes without samples (blanks) were treated similarly as sample tubes to account for background noise and other sources of contamination. Samples and blanks were sequentially derivatized as described (Misra et al 2019).

#### GC/MS Data Acquisition and Analysis

Data were generated with a high-resolution (HR) Orbitrap Mass Spectrometer (Q Exactive Orbitrap MS, Thermo Fisher) coupled to gas chromatography (GC), and details of the procedure were described by Ampong et al (2022). In all cases, 1 µL of derivatized sample was injected into the TRACE 1310 GC (Thermo Scientific, Austin, TX) in a splitless (SSL) mode at 220°C. Helium was used as a carrier gas and the flow rate was set to 1 mL/min for separation on a Thermo Scientific Trace GOLD TG-5SIL-MS (30 m length × 0.25 mm i.d. × 0.25 μm film thickness) column with an initial oven temperature of 50°C for 0.5 min, followed by an initial gradient of 20°C/min ramp rate. The final temperature of 300°C was held for 10 min. All eluting peaks were transferred through an auxiliary transfer line into a Q Exactive-GC-MS (Thermo Scientific, Bremen, Germany). Raw data obtained from data acquired by GC/MS were converted using the open source ProteoWizard’s msConvert software prior to data preprocessing with MS-DIAL 4.6 software (Riken, Japan, and Fiehn Lab, UC Davis, Davis, CA, USA). Spectral library matching for metabolite identification was performed using an in-house and public library. Data were further normalized by QC-based-loess normalization prior to log10 transformation and missing values were imputed based on random forest imputation method (Ampong et al 2022).

### Statistical Analysis of Integrated Omics Data

#### Weighted Gene Co-expression Network Analysis (WGCNA)

WGCNA was performed with the WGCNA package (Langfelder and Horvath 2008) in R software according to the R package WGCNA protocol (https://horvath.genetics.ucla.edu/html/CoexpressionNetwork/Rpackages/WGCNA/). Liver omics data from female baboons were used to create a weighted adjacency matrix by calculating Pearson correlations for pair-wise omics data. The co-omics similarity was raised to a power β = 26 to calculate adjacency (Zhang and Horvath 2005) in accordance with a scale-free network. This adjacency matrix was then transformed into a topological overlap matrix (TOM) to measure relative gene interconnectedness and proximity. The TOM was then used to calculate the corresponding dissimilarity (1 – TOM). Average linkage hierarchical clustering coupled with the TOM-based dissimilarity was used to group correlated omics data into modules (Zhang and Horvath 2005). More specifically, modules were generated from the Dynamic Tree Cut method for Branch Cutting. The major parameters were set with a deep-split value of 2 to branch splitting and a minimum size cutoff of 50 (minimum cluster size = 50) to avoid abnormal modules in the dendrogram; highly similar modules were merged with a height cutoff of 0.25. In the resulting network, as neighbors in a cluster share high topological overlap, the resulting modules likely indicate a common functional class. WGCNA has the advantage of allowing analysis of continuous traits without binning the data for arbitrary phenotypic cutoffs in an analysis, preventing loss of power due to binning.

#### Construction of Module-Trait Relationships

The omics modules summarize the main patterns of variation and the first principal component represents the summary of the module and is referred to as the module eigengene (Langfelder and Horvath 2007). Modules were associated with both age and age-squared to assess both the linear and non-linear changes with age. In addition, all modules were associated with morphometric and clinical data. The relationship between module eigengenes, morphological and clinical measures was assessed by Pearson correlation; if p-value < 0.05, then the module and clinical trait was considered significant. The modules and clinical traits that had a p-value < 0.05 and a correlation ≥ 0.30 were retained for further investigation. A heat map was used for visualization of the correlations of each module-trait relationship.

#### Pathway and Network Enrichment Analysis

To assess directionality of pathways significantly enriched with molecules from integrated omics modules, all molecules in the positively correlated modules were converted to positive fold change molecules and all molecules from negatively correlated modules were converted to negative fold change molecules. Then all molecules from positively correlated and negatively correlated significant modules were imported separately to Ingenuity Pathway Analysis (IPA) software (Qiagen) for core analysis where pathways were analyzed for significant enrichment of module genes. Pathways were ranked by -log p-value and right-tailed Fisher’s exact test was used to calculate association between molecules in the dataset and molecules in annotated pathways (Ingenuity® Systems, http://www.ingenuity.com). A p-value of < 0.01 was considered significant. We used an end-of-pathway approach to identify pathways in which activity was biologically consistent at the beginning and end of the pathway. We hypothesized that a pathway is potentially relevant if molecular changes at the end of the pathway are consistent with the overall pathway change, i.e., activated or inhibited (Nijland et al 2007).

## Results

Our study included liver samples and corresponding blood samples from 35 female baboons ranging in age from 6.0 to 22.1 y (human equivalent ∼24 to 88 y), as well as a collection of morphometric measures from these animals. Liver samples were used for transcriptomic, proteomic and metabolomic analyses, and blood samples were used to measure levels of lipoproteins and hormones related to metabolism and stress. Correlation analysis showed significant associations of age with triglycerides (p=9.5x10^-3^) and waist circumference (p=0.03), and associations of age^2^ with body length and crown rump length (p=1.0x10^-4^) (Table 1).

**Table 1.** Age or age-squared correlation with morphometric and blood clinical measures.

| Trait | Age Correlation | Age P-value | Age-squared Correlation | Age-squared P-value |
| --- | --- | --- | --- | --- |
| Total cholesterol (mg/dl) | 0.11 | 0.51 | -0.21 | 0.21 |
| HDL cholesterol (mg/dl) | -0.08 | 0.64 | -0.14 | 0.39 |
| LDL cholesterol (mg/dl) | 0.17 | 0.30 | -0.27 | 0.11 |
| Triglycerides (mg/dl) | 0.43 | **$9.5 \times 10^{-3}$** | 0.14 | 0.39 |
| Glucose (md/dl) | 0.30 | 0.07 | 0.01 | 0.91 |
| Liver cortisol (ug/dl) | 0.25 | 0.14 | 0.11 | 0.49 |
| Leptin (pg/ml) | 0.01 | 0.94 | -0.06 | 0.71 |
| Body weight (kg) | 0.13 | 0.44 | -0.15 | 0.38 |
| Body length (cm) | -0.33 | 0.05 | -0.62 | **1.0x10<sup>-4</sup>** |
| BMI (kg/m2) | 0.30 | 0.07 | 0.18 | 0.28 |
| Crown rump length (cm) | -0.19 | 0.24 | -0.63 | **1.0x10<sup>-4</sup>** |
| Abdominal circumference (cm) | 0.28 | 0.09 | 0.14 | 0.39 |
| Chest circumference (cm) | 0.10 | 0.53 | 0.08 | 0.63 |
| Waist circumference (cm) | 0.35 | **0.03** | 0.15 | 0.37 |
| Liver weight (g) | -0.02 | 0.89 | -0.21 | 0.21 |

### Integrated Omics to Identify Age-Associated Modules

Initially, we used WGCNA to identify modules of co-correlated omics molecules, and then determined whether any of these modules correlated with age and age-squared. A total of 26,311 omics molecules, consisting of 24,562 transcripts (Supplemental Tables ST2), 1427 proteins (Supplemental Tables ST3), and 322 metabolites (Supplemental Tables ST4), passed quality filters and were included in WGCNA. The results revealed 3 modules that were positively correlated with age (light yellow, p = 3.0x10^-3^, correlation = 0.49; dark red, p = 0.04, correlation = 0.35; and green, p = 0.02, correlation = 0.39) (Figure 1, Table 2, Supplemental Tables ST5, ST6), which included 3149 genes, 33 proteins, and 0 metabolites. Two modules were negatively correlated with age (sky blue, p = 9.0x10^-3^, correlation = -0.43; and white, p-value = 0.03, correlation = -0.36) (Figure 1, Table 2), which included 37 genes, 216 proteins, and 0 metabolites. WGCNA also identified one module that was negatively correlated with age-squared (brown, p = 0.04, correlation = -0.35), which included 1122 genes, 1 protein, and 0 metabolites (Figure 1, Table 2, Supplemental Tables ST6). Overall, the number of molecules in the 3 modules positively correlated with age was 12.8% of transcripts and 2.3% of proteins that passed quality filters. The number of molecules in the 2 modules negatively correlated with age was 0.15% of transcripts and 15.1% of proteins that passed quality filters.

**Figure 1:**
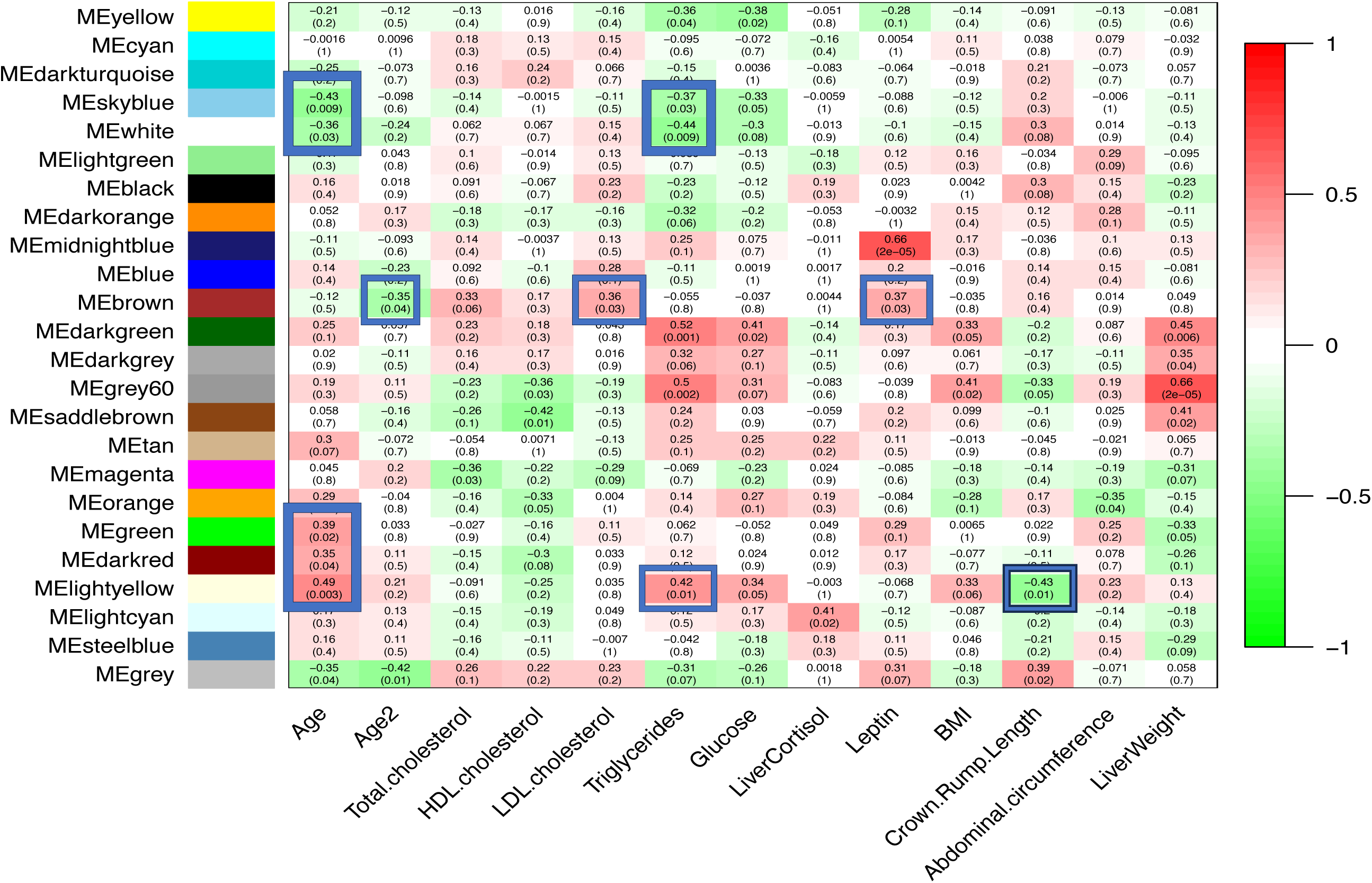
WGCNA of omics data.

**Table 2.** Number of genes, proteins, and metabolites in each significant WGCNA module correlated with age and age-squared.

| Module colors | Age |  | Age-squared |  | No. Genes per Module | No. Proteins per Module | No. Metabolites per Module |
| --- | --- | --- | --- | --- | --- | --- | --- |
|  | Pearson correlation | P-value | Pearson correlation | P-value |  |  |  |
| Sky blue | -0.434 | 0.009 | -0.098 | 0.576 | 12 | 112 | 0 |
| White | -0.359 | 0.034 | -0.236 | 0.173 | 25 | 104 | 0 |
| Brown | -0.119 | 0.496 | -0.351 | 0.039 | 1122 | 1 | 0 |
| Green | 0.395 | 0.019 | 0.033 | 0.850 | 655 | 3 | 0 |
| Dark red | 0.347 | 0.041 | 0.108 | 0.536 | 228 | 1 | 0 |
| Light yellow | 0.487 | 0.003 | 0.208 | 0.230 | 2266 | 29 | 0 |

Next, we assessed whether age and age-squared associated modules overlapped with modules that had significant associations with clinical and morphometric measures. Blood levels of triglycerides showed significant negative associations and overlapped with the age-associated sky blue and white modules (Figure 1, Supplemental Tables ST5). In addition, triglyceride showed significant positive associations and overlapped with age in the light yellow module. (Figure 1, Supplemental Tables ST5). While crown rump length was also significantly associated with the light yellow module, the association was opposite of that with age. Blood levels of LDL cholesterol and leptin were associated positively with the brown module, which was an age-squared associated module, but the association was opposite of that with age-squared (Figure 1, Supplemental Tables ST5).

### Pathway Enrichment Analysis

All molecules in the positively correlated modules were assigned positive fold changes and combined into a single list, and negatively correlated molecules were assigned negative fold changes and combined into a single list to identify enriched pathways and assess directionality in the pathways (IPA). The top five pathways identified from positively correlated modules with age are shown in Figure 2A. We focused on UPR and ER stress pathways that were positively correlated with age (Figure 3, Supplemental Tables ST7). The UPR is an adaptive response that resolves ER stress by enhancing the degradation of misfolded proteins and expanding the protein folding capacity of cells through the up-regulation of ER chaperones. The UPR showed up-regulation of ER-transmembrane transcription factors (ATF6 and ATF4), eukaryotic Initiation Factor 2α (eIF-2α), heat shock protein A5 (HSPA5), calnexin, calreticulin, DnaJ heat shock proteins, heat shock protein 90 beta family member 1 (HSP90B1) and X-box binding protein 1 (XBP1). In addition, *HSPA5,* ER-degradation enhancing α-mannosidase-like (*EDEM1*), tumor necrosis factor receptor-associated factor2 (*TRAF2),* and valosin containing protein (*VCP)* genes were observed in UPR and ER stress pathways. These genes were reported to function in many protein folding processes. The *HSPA5* gene encodes the binding immunoglobulin protein (BiP), a Hsp70 family chaperone localized in the ER lumen, that initiates UPR to decrease unfolded/misfolded proteins to maintain ER homeostasis (Wang et al 2017). The *EDEM1* gene encodes a protein that extracts misfolded proteins from the calnexin folding cycle and targets them for degradation by ER-associated protein degradation (ERAD) (Chiritoiu et al 2020). The protein encoded by the *TRAF2* gene was shown to regulate the cellular response to stress and cytokines by controlling JNK, p38 and NF-κB signaling cascades (Habelhah et al 2004), and the protein encoded by the *VCP* gene was shown to extract ubiquitinated ERAD substrates from the ER into the cytosol for proteasomal degradation (Wojcik et al 2006).

**Figure 2A:**
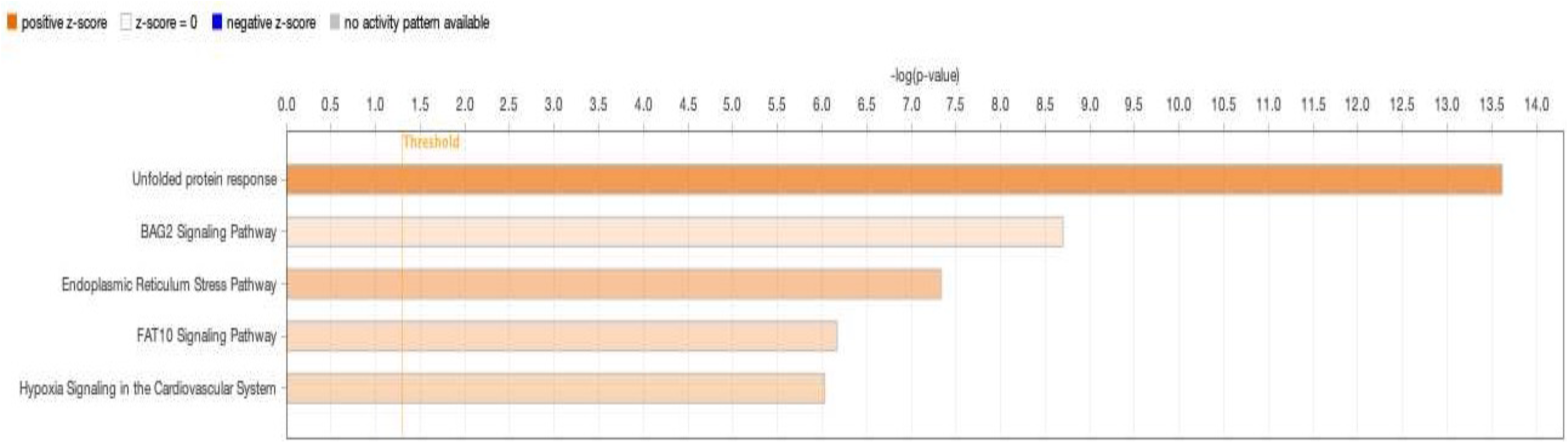
Top 5 pathways from positively correlated modules with age.

**Figure 2B:**
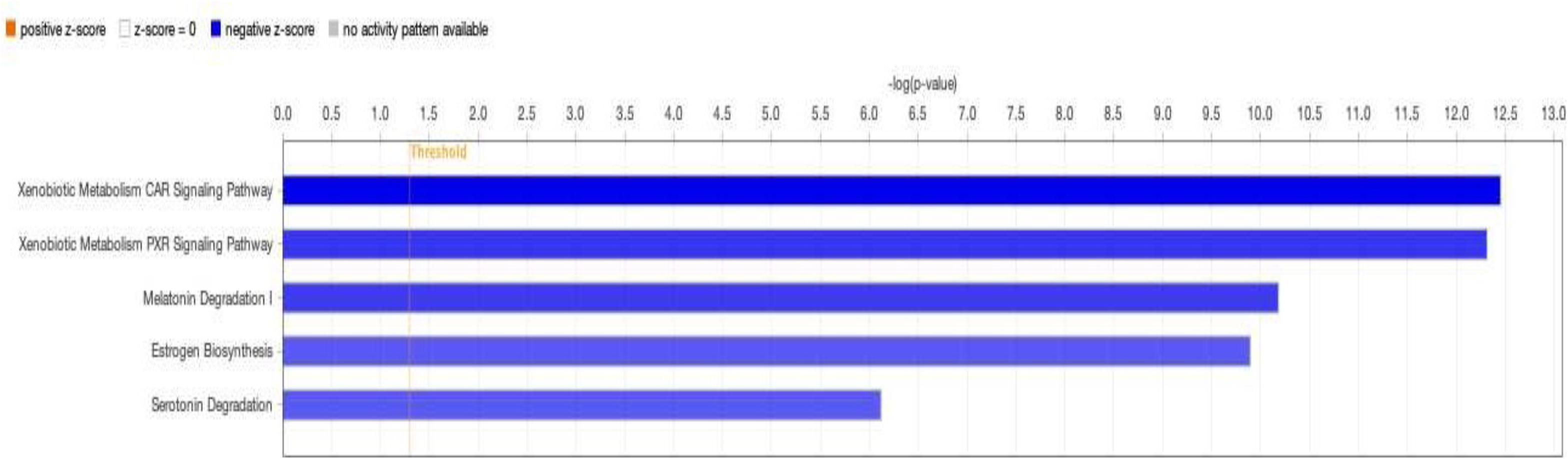
Top 5 pathways from negatively correlated modules with age.

**Figure 3:**
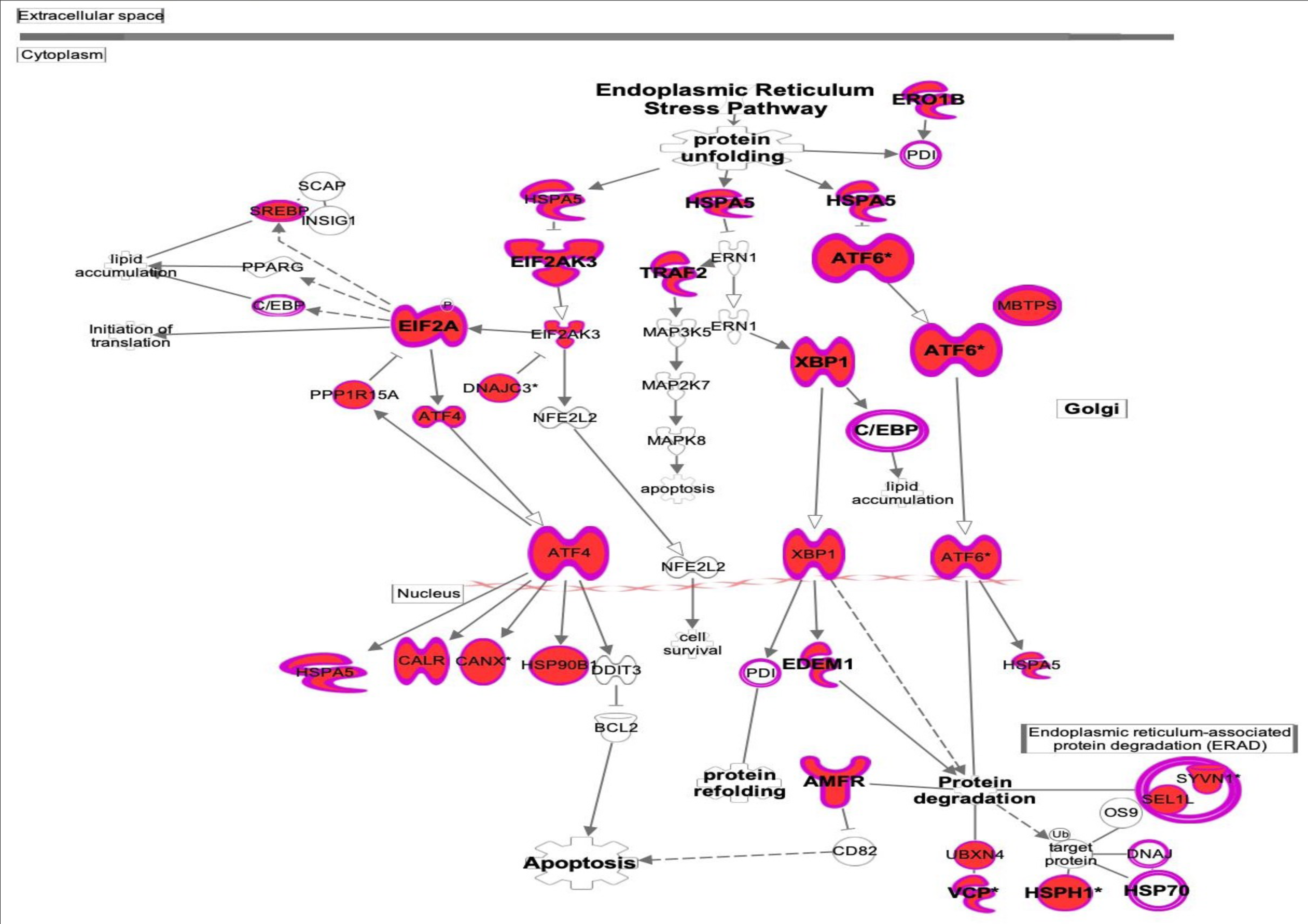
Endoplasmic reticulum stress and unfolded protein response.

The top five pathways identified from negatively correlated modules with age are shown in Figure 2B. We focused on four pathways that were negatively correlated with age and they are: xenobiotic metabolism constitutive androstane receptor (CAR) signaling, xenobiotic metabolism pregnane X receptor (PXR) signaling, melatonin degradation (Figure 4, Supplemental Tables ST8) and serotonin degradation (Figure 5, Supplemental Tables ST8).

**Figure 4:**
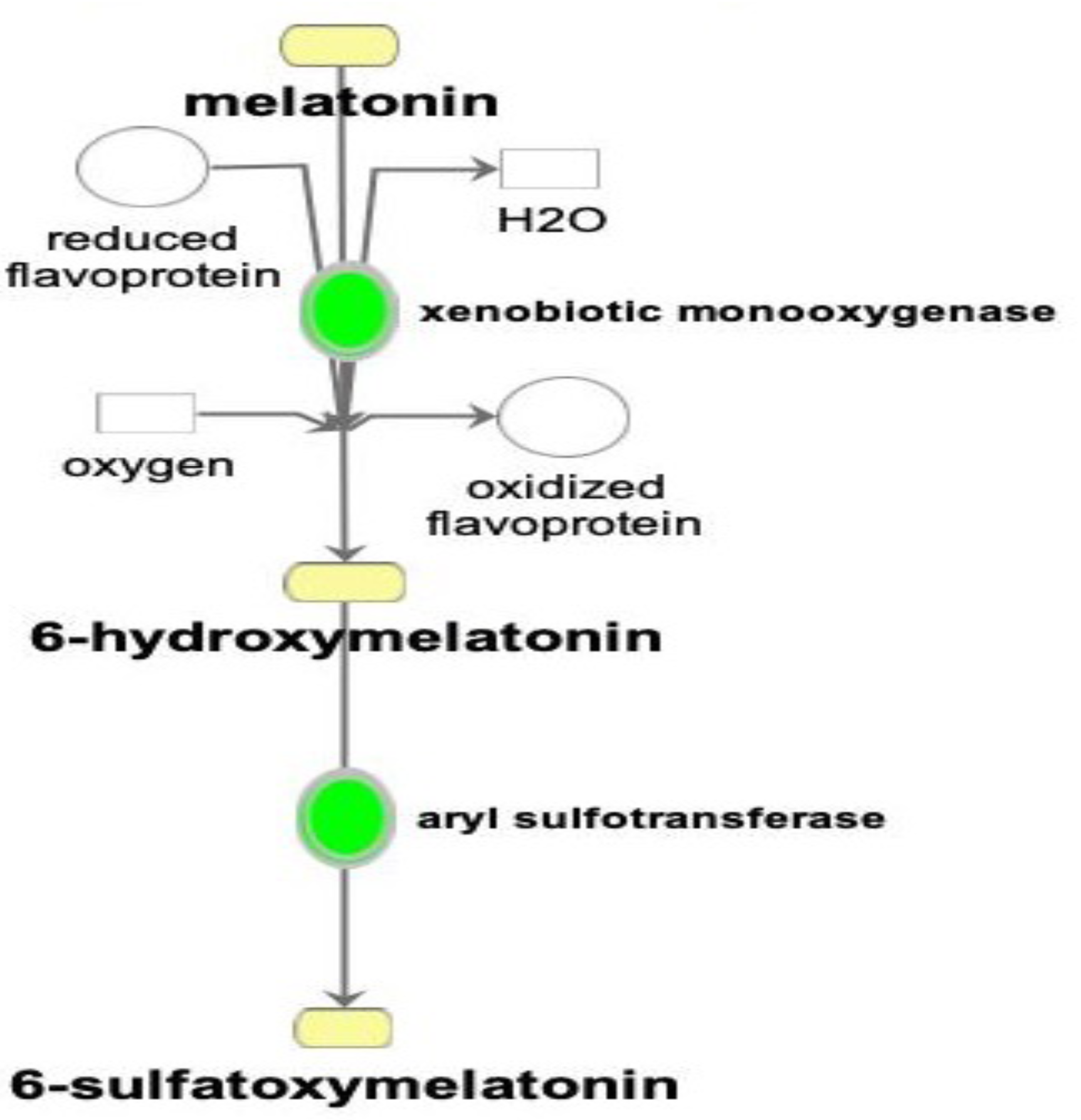
Melatonin degradation pathway.

**Figure 5:**
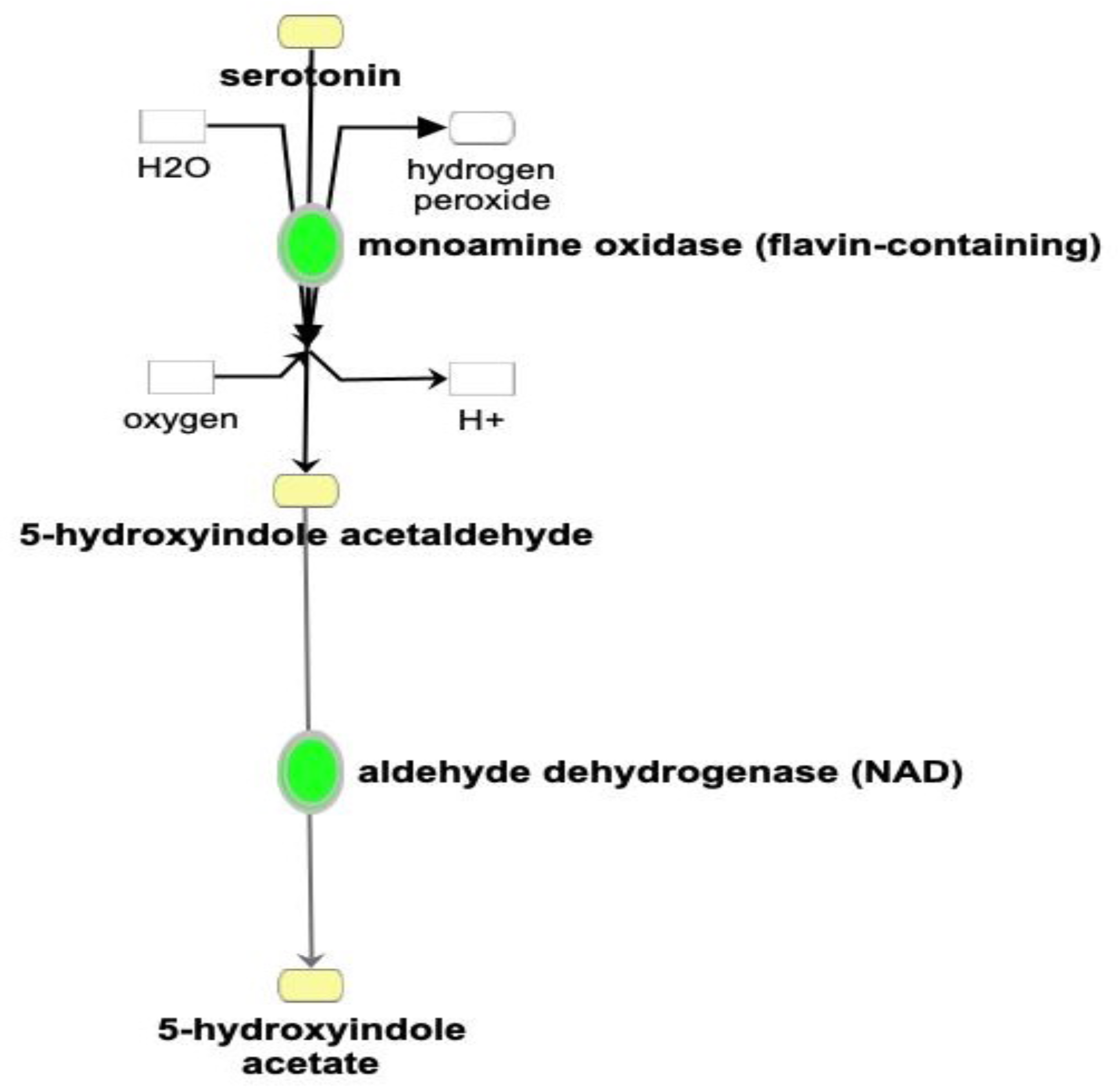
Serotonin degradation pathway.

The main pathway of melatonin degradation showed down-regulation of two enzymes, xenobiotic monooxygenase and aryl sulfotransferase (Figure 4), whose expression showed negative correlation with age. The first step in the degradation of endogenous and exogenous melatonin is mediated by cytochrome P450 superfamily of enzymes in the liver to produce 6-hydroxymelatonin (Figure 4) which is further metabolized to 6-sulfatoxymelatonin by aryl sulfotransferase in the liver and subsequently excreted in the urine (Claustrat et al 2005). The metabolite 6-hydroxymelatonin is known for its potent antioxidant property (Alvarez-Diduk et al 2015).

The degradation of serotonin involves two enzymes, monoamine oxidase A (MAO-A) and aldehyde dehydrogenase, whose expression also showed negative correlation with age (Figure 5). MAO-A is responsible for the oxidative deamination of serotonin to 5-hydroxyindole acetaldehyde (Squires et al 2006, Bortalato et al 2010), which is further metabolized to 5-hydroxyindole acetate by aldehyde dehydrogenase in the liver and subsequently excreted in the urine (Squires et al 2006). Hydrogen peroxide, a reactive oxygen species (ROS), is generated during the conversion of serotonin to 5-hydroxyindole acetaldehyde.

The xenobiotic metabolism CAR and PXR signaling pathway also showed negative correlation with age. The CAR and PXR pathways are responsible for xenobiotic detoxification and elimination to reduce oxidative stress. Expression of many enzymes, such as aldehyde dehydrogenase, cytochrome P450 enzymes, sulfotransferase, UDP-glucuronosyltransferase and glutathione s-transferase, in these two pathways was negatively correlated with age.

## Discussion

The goal of this study was to generate multi-omics data and use the WGCNA method to analyze multi-omics data together in an unbiased manner for identification of molecular pathways associated with healthy aging in the female baboon liver. It is known that aging is a risk factor in the development of many diseases, such as insulin resistance, diabetes mellitus, and NAFLD (Hunt et al 2019). A number of studies focused on molecular changes associated with liver diseases and aging due to the impact of a high-fat diet and physiological changes inherent to aging that promote the development of liver injury and inflammation, and progression to chronic liver diseases (Fontana et al 2013, Kim et al 2015) but none of the studies to date have characterized molecular changes across the adult age span in healthy NHP liver. The present study is unique because the animals in this cohort were maintained on a low cholesterol and low fat diet (healthy chow diet) throughout the adult age span approximating ∼24 to 88 y in human age.

Our multi-omics approach identified molecular changes in age-associated pathways in female baboon livers, providing evidence for molecular mechanisms associated with age. Known pathways included activation of UPR and ER stress with age. The ER plays an essential role in maintaining protein homeostasis by regulating protein synthesis, protein folding, and processing. Under conditions of ER stress, normal ER function becomes compromised leading to the accumulation of unfolded or misfolded proteins. Many types of cellular disturbances like glucose deprivation, aberrant calcium signaling, viral infection, and disruption of redox regulation provoke ER stress (Henkel and Green 2013). Upon ER stress, an adaptive cellular response, the UPR is activated to restore ER homeostasis and promote cell survival (Liu and Green 2019). Several studies showed that UPR is activated in many acute and chronic liver diseases including NAFLD and steatosis (Xia et al 2022, Liu and Green 2019, Liu et al 2015, Lake et al 2014, Henkel and Green 2013, Malhi and Kaufman 2011). Our data showed that activated ER stress and UPR increased with age, and the results imply aging could be a factor for NAFLD. The age-associated modules (in which ER stress and UPR were up-regulated) overlapped with a blood triglycerides-associated module, showing that the increase in blood triglycerides was correlated with molecular changes in ER stress and UPR in aging (Table 1). Most of the older female baboons had higher levels of circulating triglycerides than younger baboons, but the triglyceride levels of older baboons were not in the abnormal range. Therefore, even in healthy aging, ER stress seems to be a factor in elevating blood triglycerides. An elevation in blood triglycerides is a predictor of NAFLD (Xing et al 2021).

Our results showed that melatonin and serotonin degradation decreased with age. Melatonin is produced by the pineal gland, secreted into the blood, and acts as a hormone (Amaral et al 2018). Melatonin regulates many functions, including circadian rhythm, energy metabolism, and the immune system (Lee 2019, Carretero et al 2009). Previous studies showed that melatonin levels decline gradually with age (Jin et al 2021, Chrustek et al 2021). The deficiency of melatonin in the process of aging was suggested to reduce antioxidant protection in the elderly contributing to some age-related diseases (Reiter et al 1996 and 1997, Karasek 2004, Cardinali et al 2021, Gimenez et al 2022). Metabolites of melatonin (e.g., 6-hydroxymelatonin) were shown to be effective antioxidants and free radical scavengers playing an important role in resisting oxidative stress and free radical generation (add Ref). In our study, expression of melatonin degradation enzymes that produce 6-hydroxymelatonin was inversely correlated with age, suggesting that there is less antioxidant protection from 6-hydroxymelatonin with aging (Manchester et al 2015).

Serotonin is a biogenic amine that functions as a neurotransmitter with numerous functions in the central nervous system (Lesurtel et al 2012). Serotonin is also a key mediator of various biological processes in peripheral tissues and is shown to be involved in many pathological conditions of the liver (Lesurtel et al 2012). The peripheral measures of serotonin in blood tend to decrease with age (Jernej et al 1999) and serotonin also decreases with age across species (Yin et al 2014). Serotonin is mainly degraded in the liver by MAO-A and generates hydrogen peroxide (H_2_O_2_), which is a reactive oxygen species (ROS). Accumulation of ROS creates oxidative stress in cells and tissues (Pizzino et al 2017). Serotonin degradation by MAO-A was implicated as an important source of ROS in rodent models (Nocito et al 2007). In our study, MAO-A expression was inversely correlated with age, suggesting that ROS generated from serotonin decreased in older animals and would have a beneficial effect on liver health. In addition, our study showed that xenobiotic metabolism of CAR and PXR pathways decreased with age. The CAR and PXR are xenobiotic sensors predominantly expressed in the liver. The down-regulation of these pathways suggests that xenobiotic compounds are metabolized to a lesser extent, leading to accumulation of xenobiotic compounds and causing an increase in oxidative stress.

Thus, our study suggests that UPR activation and decreased degradation of serotonin with age may reduce oxidative stress in healthy baboon livers. Additionally, our study also suggests that down-regulation of the melatonin degradation pathway, and reduced xenobiotic metabolism from CAR and PXR pathways may increase oxidative stress.

## Conclusions

Our unbiased integrated omics analysis of the primate liver revealed that ER stress increased with age, and melatonin and serotonin degradation and xenobiotic metabolism of CAR and PXR signaling decreased with age. The results of our study suggest that UPR may play an important role in mitigating ER stress and reduced degradation of serotonin may afford protection from oxidative stress during the aging process. Overall, the pathways we identified in this study provide a better understanding of age-related molecular changes in a healthy liver. However, age-related changes in the ER stress response and degradation pathways with initiation of liver diseases warrant further study.

## Limitations

Although, this is the first integrated omics study involving a large NHP cohort across a wide adult age span, it included only females and potential sex-specific effects of aging on liver are yet to be determined. Several genes were identified the signaling pathways and validation of these genes is essential to confirm their biological importance in the aging liver.

## Acknowledgements

The authors would like to also acknowledge the staff at the Southwest National Primate Research Center, Texas Biomedical Research Institute for the veterinary services and animal maintenance support.

## Supplementary Material

Supplemental Table 1. Clinical and morphometric measures.

Supplemental Table 2. Normalized gene expression.

Supplemental Table 3. Normalized protein abundance.

Supplemental Table 4. Normalized metabolites.

Supplemental Table 5. WGCNA modules with significantly correlated phenotypic measures.

Supplemental Table 6. Genes, proteins, and metabolites in the age and age-squared associated modules.

Supplemental Table 7. Positive Module Pathways

Supplemental Table 8. Negative Module Pathways.

